# Epigenomic Analyses Identify FOXM1 As a Key Regulator of Anti-Tumor Immune Response in Esophageal Adenocarcinoma

**DOI:** 10.1101/2023.07.28.551066

**Authors:** Benjamin Ziman, Qian Yang, Yueyuan Zheng, Megha Sheth, Chehyun Nam, Hua Zhao, Uttam K Sinha, De-Chen Lin

**Affiliations:** Center for Craniofacial Molecular Biology, Herman Ostrow School of Dentistry, University of Southern California, 2250 Alcazar St, Los Angeles, CA 90089, USA; Department of Otolaryngology Head and Neck, Keck School of Medicine, University of Southern California, 1441 Eastlake Ave, Los Angeles, CA 90033 USA; Samuel Oschin Comprehensive Cancer Institute, Department of Medicine, Cedars-Sinai Medical Center, 127 S. San Vicente Blvd, Los Angeles, CA 90048, USA

## Abstract

Unlike most cancer types, the incidence of esophageal adenocarcinoma (EAC) has rapidly escalated in the western world over recent decades. Using whole genome bisulfite sequencing (WGBS), we identify the transcription factor (TF) FOXM1 as an important epigenetic regulator of EAC. FOXM1 plays a critical role in cellular proliferation and tumor growth in EAC patient-derived organoids and cell line models. We identify ERBB2 as an upstream regulator of the expression and transcriptional activity of FOXM1. Unexpectedly, Gene Set Enrichment Analysis (GSEA) unbiased screen reveals a prominent anti-correlation between FOXM1 and immune response pathways. Indeed, syngeneic mouse models show that FOXM1 inhibits the infiltration of CD8^+^ T cells into the tumor microenvironment. Consistently, FOXM1 suppresses CD8^+^ T cell chemotaxis *in vitro* and antigen-dependent CD8^+^ T cell killing. This study characterizes FOXM1 as a significant EAC-promoting TF and elucidates its novel function in regulating anti-tumor immune response.

## Introduction

Esophageal cancer poses a significant global health burden, ranking as the 9th most common cancer worldwide and the 6th leading cause of cancer-related deaths, with over 500,000 fatalities annually. The disease encompasses two major histological subtypes: esophageal squamous cell carcinoma (ESCC) and adenocarcinoma (EAC). Notably, the global incidence of ESCC has been declining, while EAC is on the rise, particularly in developed countries. For example, in the United States, the incidence of EAC has shown a substantial escalation, with rates soaring by over 300% between 1975 and 2004 (1). Unfortunately, EAC is often diagnosed at advanced stages, limiting treatment options, with more than half of cases being deemed unresectable (2). Conventional chemotherapies have demonstrated limited efficacy, and disease progression frequently leads to distal metastasis, resulting in an overall 5-year survival rate of a dismal 20% www.seer.cancer.gov (3).

Our mechanistic understanding of anti-tumor immune response has been markedly improved over the last 2 decades, and consequently immunotherapies such as immune checkpoint blockade (ICB) therapies have transformed the paradigm of the clinical management of many cancers, including EAC. However, only a minority of EAC patients show durable clinical responses to ICB therapies (4, 5). While incompletely understood, multiple mechanisms have been proposed to enable cancerous cells to evade immune surveillance and to resist ICB treatment. In particular, tumors exhibiting immune-cold phenotypes (limited intratumoral infiltration of immune cells) are often resistant to ICB therapies. In fact, the abundance of intratumoral T cells is among the most reliable predictors of the effectiveness of ICB treatment (6–9). However, in EAC, molecular mechanisms underlying the regulation of intratumoral trafficking of immune cells are poorly understood.

In this study, we re-analyzed our recent whole genome bisulfite sequencing (WGBS) data from EAC patient samples (10), and identified FOXM1 as a key transcription factor (TF) in the regulation of EAC epigenomics. Upregulation of FOXM1 was shown to be common in EAC tumor samples and associated with poor survival outcomes. Using EAC patient-derived organoid models and cell lines, we showed that FOXM1 regulated cellular proliferation, colony formation, and tumor growth *in vivo*. Unexpectedly, pathway enrichment analyses revealed a notable role of FOXM1 in regulating the trafficking and infiltration of T cells through the transcriptional regulation of Th1 chemokines. These data identify a significant EAC-promoting TF and elucidate a novel tumor-intrinsic function of FOXM1 in promoting immune evasion.

## Results

### DNA Methylome Sequencing Identifies FOXM1 as an important TF in EAC

We and others have previously demonstrated that loss of DNA methylation in distal regulatory elements (e.g., enhancers) represents a prominent epigenetic feature that occurs during cancer development (11–14). Leveraging this finding, we have developed a computational algorithm, ELMER (Enhancer Linking by Methylation/Expression Relationships (15), to systematically and unbiasedly identify cancer-specific TFs. Briefly, ELMER identifies cancer-specific hypomethylated regions (hypoDMRs) enriched for distal TF-binding sites (TFBS), inferred by sequence motif analysis. We applied ELMER to our recent WGBS cohort generated from 45 primary esophageal tumors and adjacent nonmalignant tissue samples (10), and identified candidate EAC-specific TFs using EAC-specific hypoDMRs (**Fig.1A**). We then inferred target genes for each candidate TF using the nearest genes of TFBS-containing hypoDMRs with open chromatin accessibility (based on ATAC-Seq peaks from TCGA EAC samples (16)). Using these predicted gene targets, we performed gene set enrichment analysis (GSEA) to compare their expression levels between EAC tumors and nonmalignant esophageal samples (**Fig.1A**). This analysis readily identified TFs that have recognized functions in EAC cancer biology, such as ETV4, HNF4A and ELF3 (**Fig.1B, C**) (14, 17, 18). Among the top ranked factors, we were particularly interested in FOXM1, a gene with important roles in other cancer types but that has not been extensively characterized in EAC (**Fig.1B, C**).

**Figure 1.**
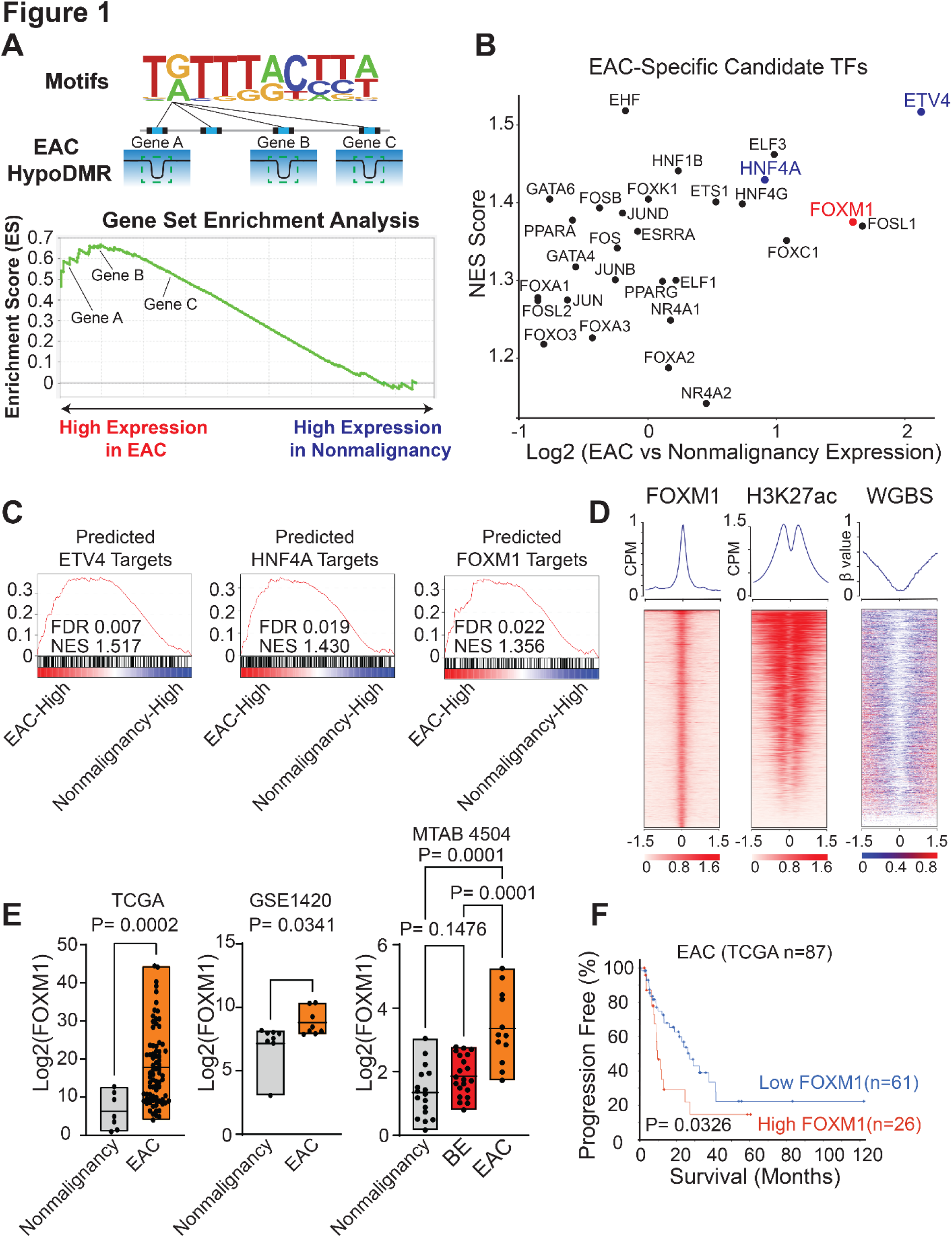
(**A)** A schematic graph of the analytic design. (**B**) A scatter plot of candidate EAC-specific TFs. The X axis represents the expression fold change between EAC and nonmalignant esophageal samples using TCGA RNA-seq data. The Y axis shows GSEA normalized enrichment score (NES) of the expression of inferred target genes of each TF. (**C**) Representative GSEA line plots for the expression of predicted target genes of ETV4, HNF4A and FOXM1. **(D)** Heatmaps of ChIP-seq signals at FOXM1 and H3K27ac peak regions and matched WGBS signal (±1.5Kb of peak center), rank ordered by intensity of peaks based on reads per million mapped reads (RPM). Lines, peaks; color scale of peak intensity shown at the bottom. **(E)** Boxplots of FOXM1 expression levels across indicated patient cohorts. P-values were determined using an unpaired t-test for 2 groups, and a one-way ANOVA with multiple comparisons for 3 groups. **(F)** Kaplan-Meier plots showing EAC patients with high FOXM1 (> mean value) and low FOXM1 expression (< mean value) from TCGA data. P-value was determined by the Log-rank test.

To validate the ELMER prediction, we first performed FOXM1 ChIP-Seq and WGBS on a matched EAC cell line (ESO26). Indeed, FOXM1 binding peaks were associated with strong depletion of DNA methylation (**Fig.1D**). Matched H3K27ac ChIP-Seq data confirmed that FOXM1 binding sites were associated with high H3K27ac signals (**Fig.1D**). Publicly available data from three different patient cohorts consistently demonstrated the upregulation of FOXM1 expression in EAC tumors compared with nonmalignant esophageal samples (**Fig.1E**). Moreover, FOXM1 expression was elevated uniquely in cancers but not in Barrett’s esophageal samples (**Fig.1E**), suggesting that its over-expression is cancer-specific rather than metaplasia-induced. High FOXM1 expression was significantly associated with poor survival outcomes of EAC patients (**Fig.1F**), indicating that FOXM1 may contribute to the biology and phenotypes of EAC.

### FOXM1 promotes malignant phenotypes of EAC

To investigate the functional role of FOXM1 in EAC, we first established a patient-derived tumor organoid model using gastroesophageal organoid culture protocols established by us and others (19, 20). We silenced FOXM1 expression using siRNAs and noted a significant reduction of organoid viability and Ki67 labeling (**Fig.2A-C**). There was also a strong decrease in organoid size following the knockdown of FOXM1 (**Fig.2D-E**). We next performed loss-of-function assays of FOXM1 using either siRNAs or shRNAs in four different EAC cell lines (**Fig.S1A-F**). Consistently, silencing of FOXM1 led to decreased proliferation of these EAC cells (**Fig.2F-G**). In addition, we performed CRISPR/Cas9 genome editing to generate single clones with FOXM1 frameshift mutations which were inducible under doxycycline (**Fig.S2G-I**). The CRISPR editing approach reproduced cellular changes observed in the FOXM1 knockdown assays (**Fig.2H**). Moreover, FOXM1 was required for colony formation of EAC cells (**Fig. 2I-J, S2A-B**). We further performed xenograft assays in immunodeficient mice and confirmed that loss of FOXM1 led to a substantial inhibition of tumor growth *in vivo* (**Fig. 2K-L**). These data demonstrate that FOXM1 promotes cellular viability and tumor growth of EAC.

**Figure 2.**
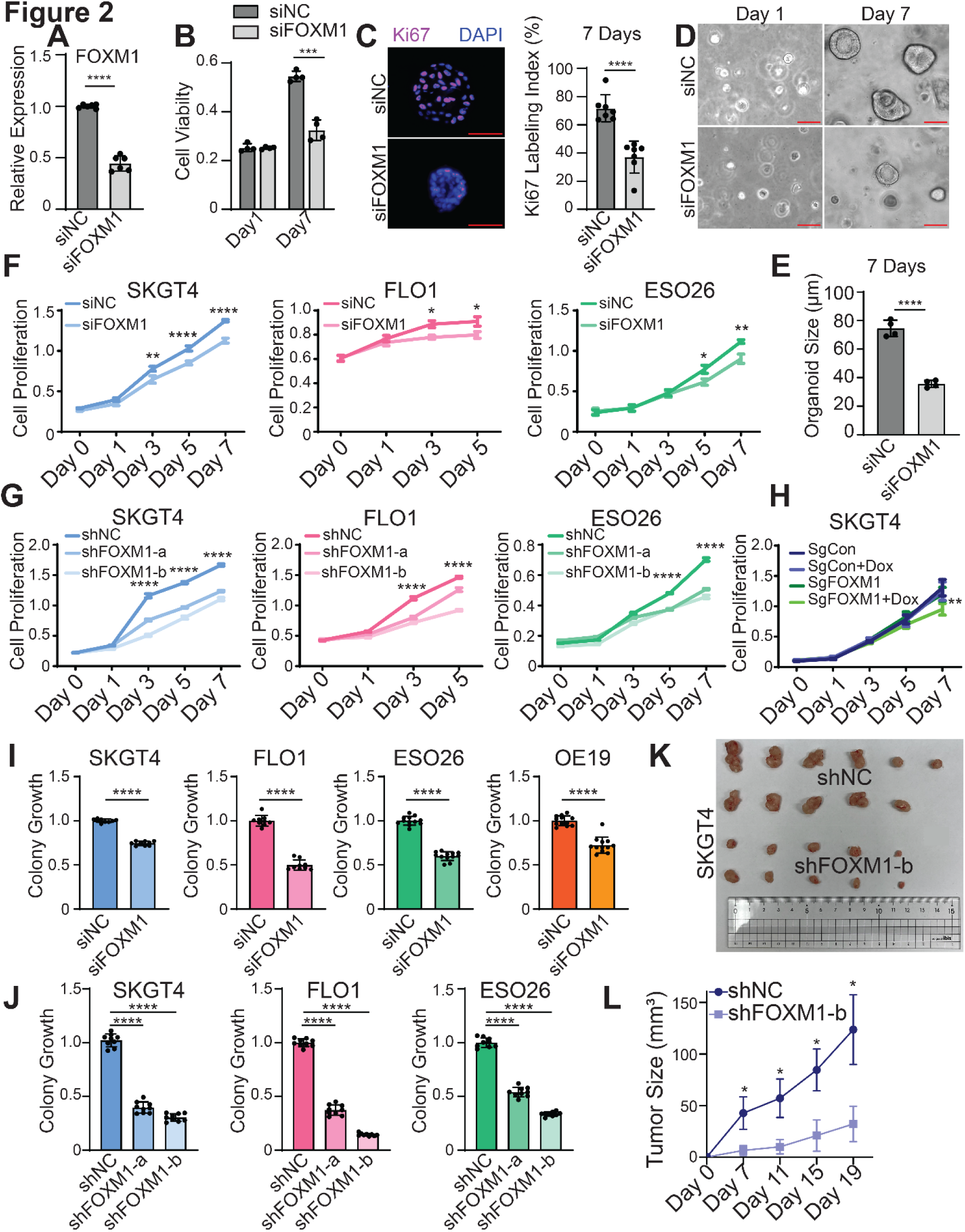
(**A**) Relative expression of FOXM1 mRNA after knockdown of FOXM1 by siRNAs in EAC patient-derived organoids. Bars show the means ± SD of two experimental replicates. ****P<0.0001. P-values were determined using an unpaired t-test. (**B**) Cell viability determined by WST-1 assay. Bars show the means ± SD of four technical replicates. ***P<0.001. P-values were determined using a multiple unpaired t-test. (**C**) Ki67 immunofluorescence staining. Scale bars, 50um. Bars show the means ± SD of seven technical replicates. ****P<0.0001; P-values were determined using an unpaired t-test (**D**) Brightfield images of representative organoids treated with siNC or siFOXM1. Scale bars, 50um. (**E**) Average organoid size determined by measuring >50 organoids. Bars show the means ± SD of four technical replicates. ****P<0.0001; P-values were determined using an unpaired t-test. **(F-H)** Cell proliferation assays in EAC cells with (**F**) FOXM1 knockdown by siRNA or (**G**) shRNAs, or (**H**) CRISPR-edited FOXM1-knockout induced by doxycycline. Graphs of **(F-H)** represent mean ± SEM of three experimental replicates. Data shown are OD values at 570nm. *P<0.05, **P<0.01, ****P<0.0001; P-values were determined using a one-way ANOVA. **(I-J)** Colony formation assays of EAC cell lines with (**I**) FOXM1 knockdown by siRNA or (**J**) shRNAs. Bar graphs of **(I-J)** represent the mean ± SD of three experimental replicates. Data are represented as fold changes relative to the control group. ****P<0.0001; P-values were determined using an unpaired t-test for 2 groups, and a one-way ANOVA with multiple comparisons for 3 groups. **(K)** Images of dissected tumor xenografts from mice. A total of 6 mice were injected in both rear flanks for each condition **(L)** Line plots of the growth of tumor size. Error bars indicate mean ± SD for each group. *P<0.05; P-values were determined using a one-way ANOVA.

### ERBB2 signaling regulates the expression and activity of FOXM1 in EAC

We recently established active enhancer profiles in EAC and showed that many oncogenic factors, such as ERBB2, are under control of EAC-specific enhancers (21). Interestingly, in our internal RNA-seq data of ESO26 cells (an ERBB2-amplified cell line), we noticed a reduction of FOXM1 mRNA upon treatment with Afatanib, an FDA-approved ERBB2-targeting inhibitor (**Fig.3A**). To formally test whether the ERBB2 signaling regulates the expression and activity of FOXM1, we began by generating a FOXM1-targeting gene signature, in order to infer FOXM1 transcriptional activity. Specifically, we first identified 252 FOXM1-occupying genes using FOXM1 ChIP-seq peaks shared by three EAC cell lines (ESO26 and SKGT4 from us; OE33 from Wiseman *et al.* (22)). We then intersected these 252 genes with down-regulated genes upon FOXM1-silencing in ESO26 cells, obtaining a 49-gene signature to predict FOXM1 transcriptional activity. Importantly, RNA-seq data following ERBB2-knockdown revealed a marked reduction of this gene signature, with 42/49 (89%) genes showing decreased expression (**Fig.3C-D**). This was accompanied by a decrease in the chromatin accessibility of the promoters of these 49 genes upon ERBB2 loss (**Fig.3E-F**). We next analyzed TCGA data from EAC patient samples, finding that ERBB2-amplified tumors had higher levels of the FOXM1 signature than tumors without ERBB2 amplification (**Fig.3G**).

**Figure 3.**
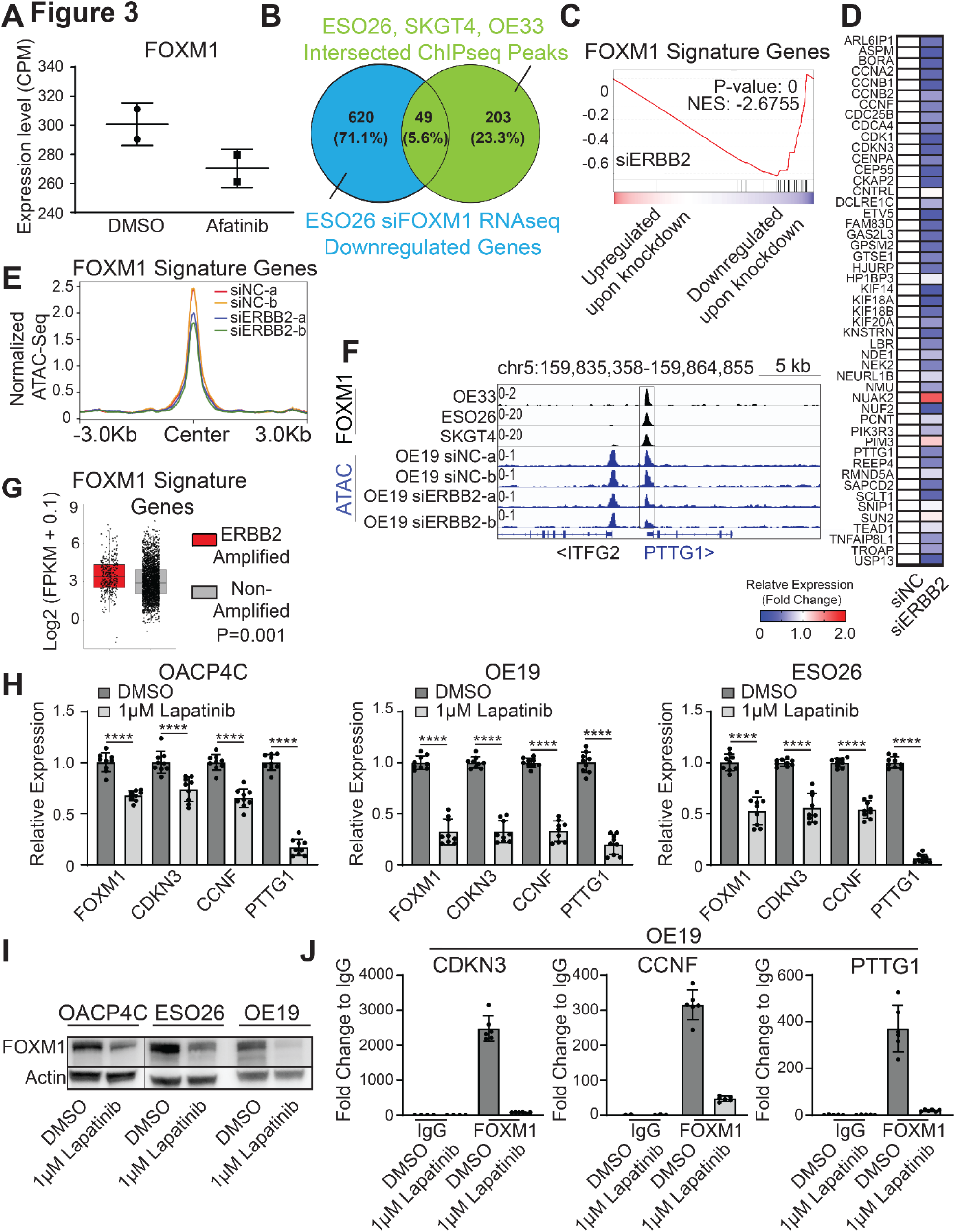
(**A**) Relative expression of FOXM1 from RNA-seq upon 24-hour treatment of 1uM Afatinib in ESO26 cells. **(B)** A Venn diagram illustrating the 49-gene intersection between the down-regulated genes from FOXM1-knockdown ESO26 cells, and the shared FOXM1 ChIP-seq peaks of 3 EAC cell lines (ESO26, SKGT4, OE33). **(C)** A GSEA line plot and (**D**) a heatmap showing the expression change of the 49-gene (FOXM1 signature) in RNA-seq upon silencing of ERBB2 in OE19 cells. In **(D),** expression changes, shown from high (red) to low (blue). Values exceeding a fold change of 2 are shown as the highest value (red). **(E)** Line plots showing the normalized ATAC-Seq signal of promoter regions for the FOXM1 signature genes following knockdown of ERBB2 in OE19 cells. **(F)** IGV plots of FOXM1 ChIP-Seq and ATAC-Seq in indicated samples. Signal values of normalized peak intensity are shown on the upper left corner. **(G)** Boxplots of the expression of 49-gene (FOXM1 signature) in EAC patients with or without ERBB2 amplifications. P-value was determined using an unpaired t-test. **(H)** Bar graphs showing relative expression of FOXM1 and its signature genes upon 24-hour treatment with 1uM lapatinib. Shown are the means ± SD from three replicates. ****P<0.0001; P-values were determined using a one-way ANOVA. **(I)** Western blotting of FOXM1 in three ERBB2-amplified EAC cell lines upon 24-hour treatment with 1uM lapatinib. Blots shown are representative of three replicates. **(J)** ChIP-qPCR assays measuring FOXM1 binding on the promoter of CDKN3, CCNF, and PTTG1 in OE19 cells. Data is represented as fold change relative to IgG control. Shown are the means ± SD with individual data points from two replicates.

After confirming the regulation of ERBB2 signaling on FOXM1 transcriptional activity, we next measured the expression of FOXM1 itself following ERBB2 inhibition. We exposed three ERBB2-amplified EAC cell lines to Lapatinib (another FDA-approved ERBB2 inhibitor), finding that Lapatinib treatment caused a decrease of FOXM1 expression at both mRNA and protein levels (**Fig.3H-I**). FOXM1 signature genes, such as CDKN3, CCNF and PTTG1, were also consistently downregulated (**Fig.3H**). We further performed ChIP-qPCR and validated that Lapatinib treatment considerably weakened the FOXM1 occupancy on its target genes (**Fig.3J**). These data identify the ERBB2 signaling as an upstream regulator of the expression and transcriptional activity of FOXM1 in EAC.

### FOXM1 inhibits immune response pathways in EAC

We next sought to reveal molecular mechanisms underlying the function of FOXM1 in EAC biology. We first performed Hallmark GSEA on RNA-Seq data following the knockdown of FOXM1 to determine pathway enrichment in EAC cells. Given that FOXM1 is a regulator of cellular proliferation, cell cycle gene sets were expectedly among the most significantly downregulated pathways following the loss of FOXM1 (**Fig.4A, left side**). Surprisingly, most of the top enriched pathways upregulated by FOXM1 knockdown belong to immune responses, including IL2-Stat5 signaling, Inflammatory responses and Allograft rejection (**Fig.4A, right side**).

**Figure 4.**
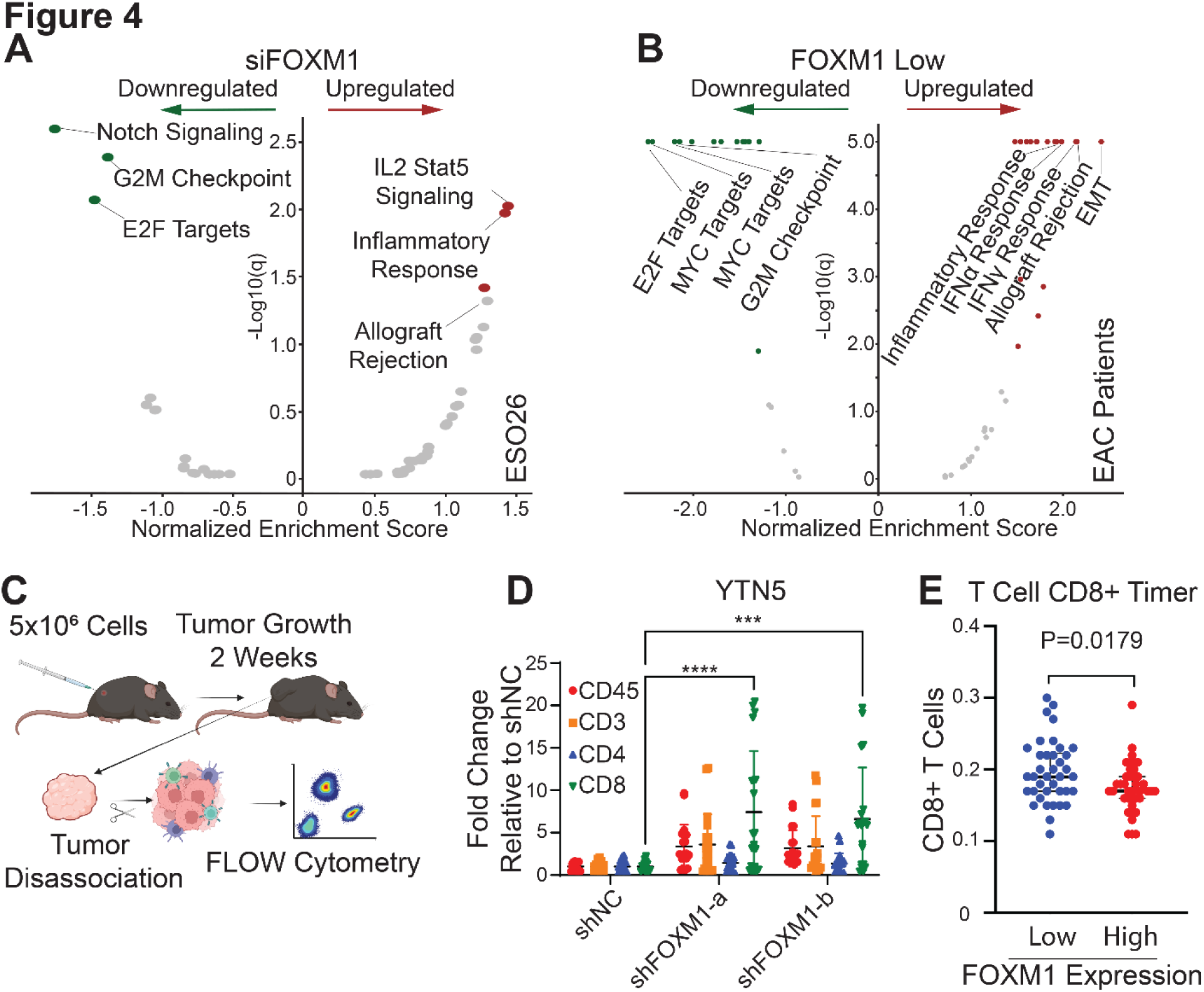
(**A-B**) Volcano plots showing GSEA results of the enriched hallmark pathways in (**A**) FOXM1-silenced ESO26 cells or (**B**) FOXM1-low EAC samples from TCGA. Significantly enriched pathways are colored. **(C)** Schematic illustration of the workflow for syngeneic xenograft experiments. **(D)** Dotplot of flow cytometry analysis of various intratumoral immune cell populations. Individual data points as well as mean ± SD from three replicates are shown. ***P<0.001, ****P<0.0001; P-values were determined using a one-way ANOVA with multiple comparisons. (**E)** Dotplots showing inferred abundance of CD8^+^ T cells in EAC tumor samples with FOXM1-low expression (bottom 50%) vs. FOXM1-high expression (top 50%). Plots show the median and interquartile range. P-value was determined using an unpaired t-test.

To corroborate these pathway enrichment results obtained from EAC cells, we next analyzed EAC patient samples and screened unbiasedly for pathways which were significantly correlated with the expression level of FOXM1 based on the TCGA RNA-Seq data. Specifically, we first stratified EAC primary samples into FOXM1-high (top 30% samples) and FOXM1-low (bottom 30% samples) groups. Next, differentially expressed genes between FOXM1-high and –low groups were used to perform GSEA (**Fig.4B**). Again, as anticipated, cell cycle pathways were enriched in the FOXM1-high group (**Fig.4B, left side**). Importantly, in the FOXM1-low group, the highest-ranking pathways were immune-related, including Allograft rejection, Interferon signaling and Inflammatory responses (**Fig.4B, right side**). Therefore, unbiased GSEA of both *in vitro* knockdown experiments and *in vivo* patient data confirmed the strong and prominent anti-correlation between FOXM1 expression and immune response pathways.

The above data prompted us to explore the functional significance of FOXM1 in regulating anti-tumor immune response *in vivo*, initially using syngeneic murine models. Because no murine EAC cell lines exist, we utilized YTN cell lines which were derived from gastric cancer models in C57BL/6 mice (23), considering that gastric cancer closely resembles EAC in both tumor biology and genomic landscapes (24, 25). We thus established xenografts in immuno-competent C57BL/6 mice using YTN cells stably expressing either scramble shRNA or FOXM1-targeting shRNAs. These tumors were collected 2 weeks post inoculation and disassociated, followed by flow cytometry analysis using markers for different immune cell populations (**Fig.4C**). Importantly, we found a significant elevation in the number of tumor-infiltrating CD8^+^ T cells in the FOXM1-knockdown group (**Fig.4D**). Consistently, deconvolution of TCGA RNA-seq data by the Timer method (26) also showed increased CD8^+^ T cells in EAC tumors with low levels of FOXM1 expression (**Fig.4E**). These data indicate that FOXM1 may inhibit T cell infiltration into the EAC tumor microenvironment.

### FOXM1 regulates the expression of Th1 chemokines

To understand how FOXM1 inhibits T cell tumor-infiltration, we focused our attention on antigen processing and presentation, as well as Th1 chemokines, which play dominating roles in T cell trafficking and recruitment (27–29). We measured the relative expression of these factors across both human EAC cells and murine gastric cancer cells. Notably, the majority of these genes were upregulated upon knockdown of FOXM1 by either siRNAs or shRNAs (**Fig.5A-B, D-E**). Consistently, ERBB2 inhibition in ERBB2-amplified cell lines elevated the expression of these immune factors (**Fig.5C**). In addition, RNA-seq data from either Afatinib-treated or ERBB2-silenced cells showed similar expression changes (**Fig.5F-G**).

**Figure 5.**
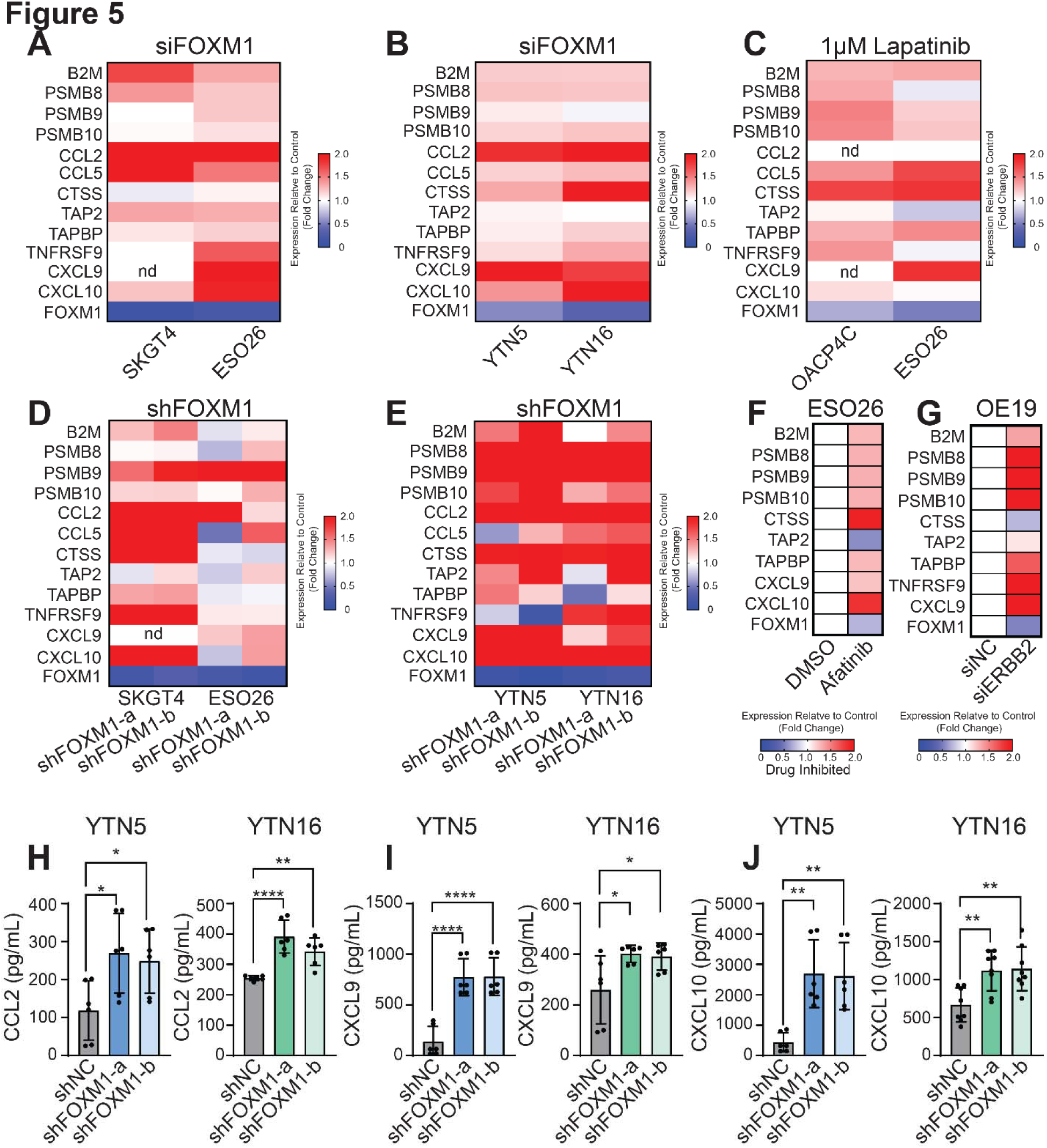
(**A-E**) Heatmaps showing the relative expression of gene expression for indicated immune factors in **(A)** EAC and **(B)** YTN cells transfected with FOXM1 siRNA, or in **(C)** ERBB2-amplified EAC cells treated with 1uM lapatinib, or **(D)** EAC and **(E)** YTN cells with stable FOXM1-knockdown. Data are shown as the geometric mean from three experimental replicates. Expression is shown from high (red) to low (blue). Values exceeding a fold change of 2 are shown as the highest value (red). Genes that were not detectable are labeled “nd”. **(F-G)** Heatmaps showing the relative expression of indicated immune genes from RNA-seq of EAC cells treated with 1uM afatinib for 24 hours **(F)** or ERBB2-knockdown **(G)**. RNA-seq did not detect the expression of CCL2, CCL5, and TNFRSF9 in ESO26 cells or CCL2, CCL5, and CXCL10 in OE19 cells. **(H)** Bar graphs showing the protein amount of secreted CCL2 (**H**), CXCL9 (**I**), and CXCL10 (**J**) in the culture media of IFNγ-stimulated YTN5 and YTN16 cells with or without FOXM1-knockdown. Bar graphs of **(H-J)** represent the mean ± SD of three experimental replicates. *P<0.05, **P<0.01, ****P<0.0001; P-values were determined using a one-way ANOVA.

To further validate the regulation of FOXM1 on the expression of these immune factors, we selected three chemokines for protein level analysis: CCL2, CXCL9, and CXCL10, given their established roles in directly recruiting T cells (27, 28, 30, 31). Indeed, ELISA assays showed that the secreted levels of these chemoattractants were significantly enhanced upon knockdown of FOXM1 (**Fig.5H-J**). These data suggest that FOXM1 regulates the expression of immune factors required for T cell recruitment and chemoattraction, particularly Th1 chemokines.

### FOXM1 regulates T cell recruitment and tumor-killing activity of CD8^+^ T cells

To establish the role of Th1 chemokines in FOXM1-regulated T cell recruitment, we performed T cell chemotaxis assays. Briefly, we incubated CD8^+^ T cells with the culture media from cancer cells in a transwell chamber, allowing for the migration of CD8^+^ T cells towards the media containing chemoattractants (**Fig.6A**). We observed that silencing of FOXM1 enhanced the migration of CD8^+^ T cells (**Fig.6B**), consistent with our *in vivo* data (**Fig.4D**). Importantly, blockade of CCL2 activity with its neutralizing antibody decreased CD8^+^ T cell migration, reversing T cell migration in the siFOXM1 group to comparable levels of the control group. We further reproduced this result by using a neutralizing antibody against CXCR3, the receptor for CXCL9 and CXCL10 on T cells (27). These results suggest that CCL2, CXCL9 and CXCL10 may act as downstream mediators of FOXM1 in recruiting CD8^+^ T cells (**Fig.5H-J, 6B**).

**Figure 6.**
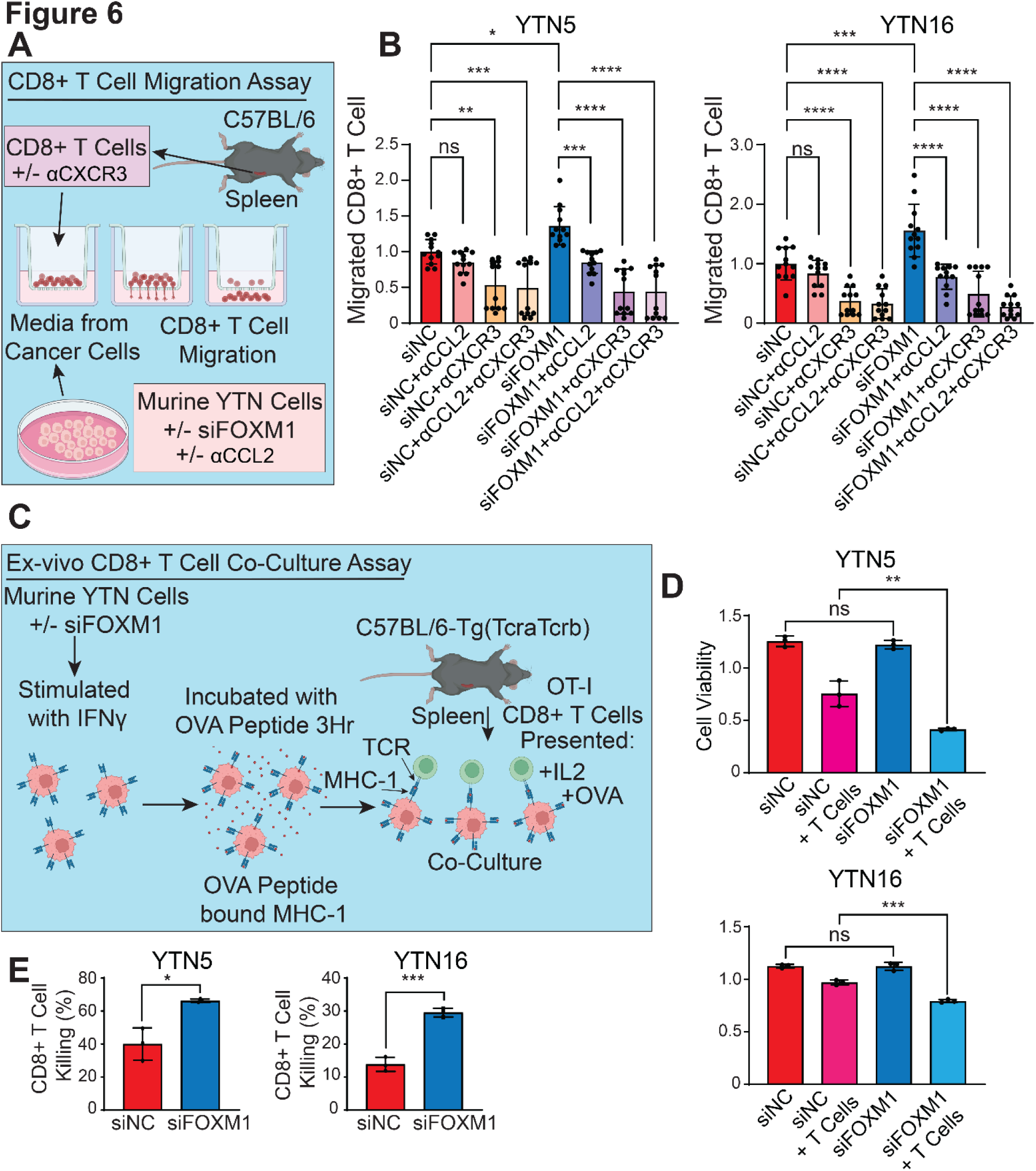
(**A**) Schematic demonstration of the workflow of CD8^+^ T cell migration assays. **(B)** Bar graphs showing the relative number of migrated CD8^+^ T cells using culture media from either YTN5 or YTN16 cells. Bars show means ± SD from two replicates. *P<0.05, **P<0.01, ***P<0.001, ****P<0.0001; P-values were determined using a one-way ANOVA with multiple comparisons. (**C**) Schematic demonstration of the workflow of *ex vivo* CD8^+^ T cell co-culture assay. **(D)** Bar graphs showing relative cell viability of YTN5 and YTN16 cells co-cultured with CD8^+^ T cells as described in **(C)**. Bars show the means ± SD representative of two replicates. **P<0.01, ***P<0.001; P-values were determined using a one-way ANOVA with multiple comparisons. **(E)** Bar graphs showing CD8^+^ T cell cytotoxic killing of YTN5 and YTN16 cells as described in **(C)**. Bars show the means ± SD representative from two replicates. Data is represented as the ratio of non-viable cells in the presence of CD8^+^ T cells, divided by the viable cells in the absence of CD8^+^ T cells from **(D)**. *P<0.05, ***P<0.001; P-values were determined using an unpaired t-test.

The above data prompted us to further test whether the increased T cell infiltration by FOXM1-silencing can lead to enhanced tumor-killing by CD8^+^ T cells. We established an *ex vivo* co-culture system by pulsing YTN cells with ovalbumin (OVA) peptide (SIINFEKL), which binds to cell surface MHC-I (32). YTN cells were then washed to remove unbound peptide before co-culture with TCR transgenic OT-I CD8^+^ T cells that specifically recognize MHC-I-bound OVA peptide (**Fig.6C**). This assay thus creates a specific antigen-dependent killing of cancer cells by CD8^+^ T cells. We found that YTN cells were largely insensitive to antigen-specific T cell killing and 60-85% tumor cells remained alive at 4:1 co-culture ratio (**Fig.6D-E**). Importantly, this resistance was significantly reversed by FOXM1-knockdown in two different YTN cells (**Fig.6D-E**). These data establish the functional importance of FOXM1 in inhibiting the CD8^+^ T cell tumor-infiltration and tumor-killing cytotoxicity against EAC cells.

## Discussion

The present study provides evidence for the functional significance of FOXM1 in EAC biology, through integrative epigenomic analyses of WGBS, ChIP-seq, RNA-seq, and functional assays. We have elucidated the role of FOXM1 in regulating cellular proliferation and tumor growth in EAC patient-derived organoids and cell lines and identified the ERBB2 signaling as an upstream regulator of FOXM1. Unbiased GSEA screen shows that immune-related signaling are among the most enriched pathways upregulated in FOXM1-silenced EAC cells and FOXM1-low patient samples. Indeed, we find that FOXM1 inhibits CD8^+^ T cell tumor infiltration, *in vitro* T cell chemotaxis, and *ex vivo* CD8^+^ T cell killing. At the molecular level, FOXM1 regulates the expression of Th1 chemokines, which are critical for the intratumoral recruitment of CD8^+^ T cells.

Increased expression of FOXM1 in cancer has been observed in various tumor types (33–35). However, the mechanistic regulation of its cancer-specific overexpression has not been well characterized. In this study, we first constructed a high-confident FOXM1-targeting gene signature by integrating RNA-seq and ChIP-seq data. Using this gene signature, we found that inhibition of ERBB2 signaling resulted in reduced FOXM1 epigenetic activity. We further showed that the ERBB2 signaling promoted the expression of FOXM1 itself. These findings suggest a crosstalk between the ERBB2 signaling and FOXM1 activity in EAC. ERBB2 is one of the most frequently amplified oncogenes (∼32%) in EAC patients (36). Notably, the monoclonal anti-HER2 antibody trastuzumab is the only FDA-approved gene targeted therapy for metastatic EAC (36–38). Given the prominent clinical significance of ERBB2, the molecular mechanism identified in this study linking FOXM1 to ERBB2 signaling has both translational and basic implications for EAC research.

FOXM1 has been recognized to play a crucial role in cell cycle transition. For example, in gastric cancer, upregulation of FOXM1 contributes to disease progression by regulating key processes such as cell cycle and epithelial-mesenchymal transition (EMT) signaling (39). In EAC, FOXM1 expression has been reported to be elevated in tumors compared to normal tissues, and to drive cell cycle gene expression (22, 40). Nonetheless, the functional contribution of FOXM1 beyond cell cycle regulation has been incompletely understood.

Here, we revealed an unexpected association between FOXM1 and immune response pathways. Unbiased GSEA analysis showed that immune-related pathways were enriched in EAC cells with knockdown of FOXM1. Consistent with this finding, EAC patient tumors with low levels of FOXM1 expression exhibited enriched immune response pathways as well as increased intratumoral CD8^+^ T cell levels. Using syngeneic murine models, we found that loss of FOXM1 resulted in heightened infiltration of CD8^+^ T cells into the tumor microenvironment, which was corroborated by *in vitro* CD8^+^ T cell chemotaxis assays. Interestingly, a recent study showed that upon inhibition of FOXM1 using Thiostrepton in Lewis lung carcinoma cells caused an increase in the number of intratumoral CD3^+^ T cells (41), in line with our observations. *Ex vivo* co-culture assays revealed that FOXM1 inhibited CD8^+^ T cell-mediated antigen-dependent killing of cancer cells. Mechanistically, we identified that loss of FOXM1 increased the expression of genes involved in antigen processing and presentation, as well as secreted chemoattractants important for CD8^+^ T cell recruitment, including CCL2 and CXCL9/CXCL10. These results together highlight the potential role of FOXM1 in modulating anti-tumor immune response.

## Methods

### Human and mouse cell lines

Human esophageal cancer cells lines OE19, ESO26, OACP4C and SKGT4 cells were grown in RPMI-1640 medium (Corning, 45000-396). Flo-1 cells were cultured in Dulbecco’s modified Eagle medium (DMEM) (Corning, 45000-304). Both media were supplemented with 10% FBS (Omega Scientific, FB-02) and 1% penicillin-streptomycin sulfate (Gibco, 10378016). Murine gastric cancer cells, YTN 5 and YTN16 were kindly provided by Dr. Sachiyo Nomura at The University of Tokyo. YTN5 and YTN16 cells were cultured in DMEM (Corning, 45000-304), with 10% heat-inactivated fetal bovine serum (Omega Scientific, FB-02) and 1% penicillin-streptomycin sulfate (Gibco, 10378016), 1X GlutaMAX (Gibco, 35050-061) and MITO+ serum extender (Corning, 355006). All cultures were maintained in a 37°C incubator supplemented with 5% CO2. All the cell lines were tested for mycoplasma and verified by us using short tandem repeat analysis.

### Patient Samples

In accordance with the approved Institutional Review Board protocols at USC, primary human EAC tumor samples were obtained from patients undergoing surgical procedures under written informed consent. Tissue samples used to generate organoids were pathologically confirmed as EAC.

### EAC patient-derived organoid culture

Fresh EAC tumor samples were transferred into ice cold conditioned PBS (formula: 10uM rho-associated kinase (ROCK) inhibitor Y27632 (Sigma, #Y0503), 2% penicillin-streptomycin sulfate (Gibco, #10378016), and 1X Primocin (InvivoGen, #ant-pm-1)). Biopsies were washed more than five times with conditioned PBS and then minced using micro-dissecting scissors into fragments <1mm3. Dissected tissue was then dissociated in digestion buffer (formula: DMEM/F12 (Corning, #10-090-CVR) containing 1% (v/v) penicillin/streptomycin, collagenase type IX (1 mg/ml) (Sigma, #C7657), and dispase type II (120 μg/ml) (Thermo Fisher, # 17105041) at 37°C for 90 minutes rocking at 200 rpm. Samples were then centrifuged at 400 × g for 3 min. Cell pellets were resuspended in 70% basement membrane extract (BME) (R&D Systems, #3533-010-02). Each droplet of cell clusters contained roughly ∼2,000 cells/ 50uL. Droplets were placed (3/well) into 24-well plates and incubated at 37°C for 10 minutes. Following the incubation, 500uL of culture medium was added to each well (formula: DMEM/F12+ supplemented with 50% Wnt-3a conditioned medium (homemade), 20% R-spondin-1 conditioned medium (homemade), 10 nM PGE2 (Sigma, #P0409), 10 ng/ml FGF-10 (Pepro Tech, #100-26), 50 ng/ml rEGF (Sigma, #E9644), 100 ng/ml Noggin (Pepro Tech, #250-38), 1mM N-acetylcysteine (Sigma, #A9165), 10 mM Nicotinamide (Sigma, #N0636), 10 nM Gastrin I (Sigma, #SCP0150), 500 nM A-83-01 (Sigma, #SML0788), 10 μM SB202190 (Sigma, #S7067), 10 μM Y27632, 1 × Primocin, 1 × B27 (Thermo Fisher, #17504044), and 1x P/S). Culture medium was replaced every 3 days. Organoids were digested with TrypLE (Thermo Fisher, #12604013) for 5 to 7 min at 37°C for passage. DMEM/F12 was then added to stop the digestion process and organoids were mechanically digested using a pipet. Samples were then centrifuged at 500 × g for 3 min at 4 °C. Supernatant was removed and the cell clusters were resuspended in 70% BME.

### Electroporation of siRNAs into organoids

Organoids were dissociated into clusters of 5 to 10 cells, resuspended in 100 μl of Opti-MEM and mixed with 5 μl of 100 μM electroporation enhancer and 10 μl of 50 μM of siRNA (Horizon Discovery, #001810-10-05 and #009762-00-0005). The mixture was carefully transferred into a precooled 2-mm electroporation cuvette. Electroporation of the organoids was performed using the NEPA21 system (Nepa Gene), with the same parameters as previously described (19). Immediately after electroporation, 400 μl of prewarmed EAC culture medium was added to the electroporation cuvette. The cells were then incubated at 37°C for 40 minutes before reseeding.

### WST-1 assay of organoids

To quantify metabolically active viable cells, EAC organoids were seeded onto 48-well culture plates. After culture for indicated intervals, 10 μl per 100 μl culture medium of WST-1 reagent (Cell Biolabs, CBA-253) were added to each well, followed by a 90-minute incubation at 37℃, 5% CO2. Following the incubation, only the medium was transferred to the wells of a new 96-well plate, and the absorbance at 450 nm was measured using a microplate reader.

### Immunofluorescence staining of organoids

EAC organoid cultures were fixed in 4% paraformaldehyde (Electron Microscopy Sciences, 15711) for 30 min. After washing in PBS for three times, organoids were embedded in 2% agarose, dehydrated for making paraffin blocks, and sectioned into 5-μm slices. Slides were deparaffinized and rehydrated, followed by antigen retrieval in sub-boiling 10 mM sodium citrate buffer (pH 6.0) for 10 min. Slides were permeabilized in 0.3% Triton X-100 in PBS and blocked in 1% BSA in PBS for 30 min at room temperature. After blocking, slides were incubated with anti-Ki67 (1:1000) (Abcam, #16667) for 3 hours in a humidified chamber at room temperature. Sections were washed by PBS with 0.1% Tween 20 (PBST) (three times for 5 min each) and incubated with Alexa Fluor secondary antibodies (1:500) (Thermofisher, #A-11036) for 1 hour. After washing with PBST, slides were mounted with Fluoroshield with DAPI (Sigma, #SLCD7376). Images were acquired with Keyence BZ-X810 microscopy.

### Cell proliferation and colony formation assays

Cells were seeded in 100uL of media into 96-well plates (2,000–8,000 cells/well) in quadruplicates. Cell proliferation was measured using 10uL of MTT (3-(4, 5-dimethylthiazol-2-yl)-2, 5-diphenyl tetrazolium bromide) (Sigma, #475989) staining for 3-hours. Crystals were dissolved by adding 100uL of 10% SDS, 0.01M HCl and incubating plates overnight. Absorbance readings were taken using a 570nm wavelength. For colony formation assay, cells were seeded into six-well plates (1,000– 4,000 cells/well) and cultured for 2–3 weeks. Colonies were fixed using 4% paraformaldehyde (Electron Microscopy Sciences, 15711) and stained using 1% crystal violet (Sigma, V5265). Colonies were then dissolved by adding 100uL of 10% SDS while shaking at room temperature for 15-minutes and absorbance readings were taken using a 595nm wavelength.

### RNA extraction, cDNA synthesis, and quantitative PCR

Total RNA was extracted using RNeasy Mini kit (QIAGEN, 70106) and cDNA was obtained from the total RNA using Maxima™ H Minus cDNA Synthesis Master Mix with dsDNase (Thermo Scientific, M1682). Quantitative PCR (qRT-PCR) was conducted with PowerUp™ SYBR™ Green Master Mix (Thermo Scientific, A25918). TATABP was used for normalization. Primers used in this study were listed in Supplementary Table 1.

### Chromatin immunoprecipitation (ChIP)

ChIP assay was performed as previously described (42). Briefly, cells were cultured ∼1 × 10^7^ cells in 10 cm dishes. Cells were fixed in 10mL of fresh media containing 1% paraformaldehyde for 10 minutes at room temperature. Cells were then quenched with 1ml of 1.25M glycine for no more than 5 minutes, followed by three washes with ice cold 1X PBS. Cells were scraped off and transferred to 15mL conical tubes. Cells were spun down at 1,500 x g for 5 minutes and lysed twice with 1 ml lysis buffer (formula:150mM NaCl, 5 mM EDTA pH 7.5, 50mM Tris pH 7.5, 0.5% NP-40) containing protease inhibitors (Roche, 04693124001). Cells were first lysed mechanically by pipetting up and down several times in a microcentrifuge tube and incubated on ice for 5 minutes. Next, cells were passed through a 23G/1mL syringe, up and down five times, and further incubated on ice for 5 minutes. Cells were then centrifuged at 9,400 × g for 5 minutes at 4 °C and the supernatant was discarded. Another 1mL of lysis buffer was added to the cells and the lysis process was repeated. Next, cell pellets were resuspended in 1mL shearing buffer (formula: 1% SDS, 10mM EDTA pH 8.0, 50 mM Tris pH 8.0) and were sonicated using a Covaris sonicator. Following sonication, samples were then centrifuged at 20,000 × g for 10 min at 4 °C and the supernatants were collected. A 10uL aliquot of the supernatants was set aside and stored at –20C to be used for the input, and the remaining supernatant was further diluted 1:5 in dilution buffer (formula: 1.1% Triton X-100, 0.1% SDS, 1.2mM EDTA pH 8.0, 167 mM NaCl, 16.7mM Tris pH 8.0). 2 μg FOXM1 antibody (Cell Signaling, 20459) was then added and incubated at 4 °C overnight on a rotating platform. The following day, 30uL of Dynabeads Protein G beads (Thermo Scientific, 10004D) were added to the samples and incubated at 4 °C for 4 hours on a rotating platform. These Dynabeads were separated by a magnet and were washed eight times with ice cold lysis buffer and twice with ice cold TE buffer (formula: 10mM Tris pH 8.0, 1 mM EDTA pH 7.5, adjusted to a final pH of 7.6). DNA samples were released by adding 100uL of reverse crosslinking buffer (formula: 136mM NaHCO3, 0.96% SDS) twice for 15 minutes while rocking at room temperature. The supernatant was then pooled together and combined with 4uL of 5M NaCl and incubated at 65 °C for 14-16 hours. Next, 1uL of RNase (Thermos Scientific, # EN0531) was added to samples and incubated at 37 °C for 30 minutes, followed by 4uL 0.5M EDTA pH 8.0 with 8uL of 1M Tris pH 7.0, and 1uL of Proteinase K (10mg/mL) (Invitrogen, #100005393) at 45°C for 1 hour. Samples were then purified by 500uL of Buffer PB (Qiagen, 19066) and vortexing, followed by transferring to a spin column and incubating for 2 minutes at room temperature. Samples were washed by Buffer PE (Qiagen, 19065) three times, and DNA was eluted by adding 55uL of sterile water to the columns and incubated for 5 minutes before being spun at 17,900g for 2 minutes. The final products were either subjected to DNA library preparation and deep sequencing using Illumina HiSeq platform or used for quantitative PCR.

### Western blotting

Cells were lysed using the RIPA lysis buffer system (Santa Cruz Biotechnologies, sc-24948A), supplemented with proteinase inhibitor cocktail and PMSF for 30 min on ice. Protein concentrations were determined with Bradford reagent (VWR, E530-1L). Western blotting was performed as previously described (43) using SDS-PAGE gels (GenScript, #M41210) and transferred to 0.45um immune-blot PVDF membrane (Millipore, #IPVH00010) for 1 hour at 100V using Transfer buffer (formula: 25mM Tris, 192mM Glycine, 20% Methanol). Membranes were blocked and primary antibodies were diluted in TBST (formula: 20mM Tris, pH 7.4, 140mM NaCl, 0.1% Tween-20) with 5% BSA (Research Products International, #9048-46-8). Primary antibodies were incubated at 4°C for 14-16 hours. Membranes were washed 4 times for 5 minutes with TBST and then secondary antibodies were incubated for 1 hour at room temperature in TBST-SDS (formula: 20mM Tris, pH 7.4, 140mM NaCl, 0.2% Tween-20, 0.01% SDS) with 5% non-fat powdered milk (Bioworld, #30620074-1). Membranes were then washed 4 times for 5 minutes with TBST and twice with 1X PBS. Membranes were developed with an ECL substrate (Thermo Scientific, A38554) and chemiluminescence. Primary antibodies used were FOXM1 (1:1,000) (Cell Signaling Technology, 20459) and Actin (1:3,000) (Developmental Studies Hybridoma Bank, JLA20). Secondary antibodies were HRP, goat anti-rabbit (Millipore Sigma, 112-348) 1:1,000 for FOXM1 and HRP, goat anti-mouse (Invitrogen, G21040) 1:5,000 for Actin.

### Transfections and lentiviral production

Cells were transfected with non-silencing siRNA or siFOXM1 (Horizon Discovery, #001810-10-05, #057933-01-0005 & 009762-00-0005) using Lipofectomine RNAiMAX (Thermo Fisher, #13778150). Lentiviral cloning vector pLKO.1-TRC (Addgene, #10878) was used for shRNA expression. The double-stranded oligonucleotide shRNAs for mouse FOXM1 sequences were ligated into the AgeI/EcoRI sites of the pLKO.1-TRC digested lentiviral vector. shRNA target sequences were listed in Supplementary Table 2. Human shFOXM1 lentiviral vectors were provided as a gift from H Phillip Koeffler’s lab. Transfection of second-generation lentiviral vectors included 2ug of lentiviral vector (pLKO.1 TRC), packaging vectors pMD2.G 0.5ug (Addgene, #12259) and pPAX2 1.5ug (Addgene, #12260), using 100uL of serum free DMEM media (Corning, 45000-304) and 6uL of BioT lipofectamine (Bioland Scientific, #B01-00). For CRISPR/Cas9 cloning, the double-stranded oligonucleotide sgRNAs for human FOXM1 sequences were ligated into the BbsI sites of the FgH1tUTG plasmid (Addgene, #70183). sgRNA target sequences were listed in Supplementary Table 2. Transfection of third generation lentiviral vectors included 2ug of lentiviral vectors FgH1tUTG and FUCas9Cherry (Addgene, #70182), packaging vectors pRSV-Rev 1ug (Addgene, #12253), pMDLg/pRRE 1.5ug (Addgene, #12251), and pMD2.G 0.5ug (Addgene, #12259) using 100uL of serum free DMEM media (Corning, 45000-304) and 6uL of BioT lipofectamine. To produce viral particles, the recombinant viral vectors and packaging vectors were co-transfected into HEK293T cells. Supernatants were harvested and filtered through a 0.45 μM filter 48 hours after transfection. Cells were then infected with the virus in the presence of 5 ug/ml Polybrene (Santa Cruz Biotechnology, #134220). All sequences were verified using Sanger sequencing (Laragen, USA).

### Selection of stable knockdown and knockout cells

Upon infection of EAC and mouse YTN cells with pLKO.1-TRC viral particles, cells were selected using puromycin (Santa Cruz Biotechnology, #205821). ESO26, FLO1, and SKGT4 cells required 3ug/mL, 4ug/mL, and 5ug/mL of puromycin, respectively. YTN5 and YTN16 cells required 8ug/mL of puromycin. Stable knockdown cells remained in media containing puromycin throughout the length of culture, and knockdown was confirmed using quantitative PCR prior to experiments. CRISPR/Cas9 viral particles were exposed to SKGT4 cells. mCherry+ and GFP+ cells were sorted, and single cells were seeded into 96 well plates. Single colony cells were maintained in culture media with tetracycline-free FBS (Omega Scientific, #FB-15). FOXM1 mutation was induced with the addition of 2.5ug/mL doxycycline (Santa Cruz Biotechnology, #100929-47-3) for 1 week.

### Xenograft studies

Animal studies were performed in accordance with protocols approved by the ethical regulations of Institutional Animal Care and Use Committee (IACUC) of USC. Xenograft models were established by subcutaneous injecting cells mixed 1:1 with Matrigel (Corning, #356238) into the rear flanks of mice. For syngeneic studies, 5 × 10^6^ YTN5 Scramble or shFOXM1 cells were injected into C57BL/6 mice (six weeks old) and tumors were allowed to grow for 2 weeks, at which point they were removed for further studies. For EAC studies, 2 × 10^6^ SKGT4 scramble or shFOXM1 cells were injected into NOD-SCID Gamma (NSG) mice (six weeks old). Mice general behaviors were monitored, and the tumor size was measured every 4 days beginning at day 7 for a total of 19 days. At the end of the experiments, mice were sacrificed, and the tumor tissues were collected for growth analysis.

### Flow Cytometry

Tumors from C57BL/6 mice were digested with collagenase/Hyaluronidase (Stemcell Technologies, #07912) and DNase1 solution (Stemcell Technologies, #07900) in DMEM media (Corning, 45000-304) to obtain single cell suspensions. Single cells (1 x 10^6^) were incubated with FC blocker (BD, #553141) at 4°C for 10 minutes. Cells were then stained with the following antibodies: Zombie Violet (BioLegend, #423113), CD3 conjugated PerCP/Cyanine (BioLegend, #100217), CD4 conjugated FITC (BioLegend, #100405), CD8 conjugated APC (BioLegend, #100711), CD45 conjugated BV605 (BioLegend, #103155) at 4°C for 30 minutes in the dark. Cells were spun down at 1,500 x g for 5 minutes and resuspended in 1mL of 1X PBS. Flow cytometry analysis was performed using the AttuneNXT machine. Populations of CD45^+^CD3^+^CD4^+^ and CD45^+^CD3^+^CD8^+^ T cells were counted and analyzed.

### Trans-well migration assay

In a 24 well plate, 400uL of culture media was collected from YTN5 and YTN16 cells first transfected with FOXM1 siRNA for 24 hours, exposure to 20ng/mL IFNγ (Abcam, #9922) for 24 hours, followed by 24-hour treatment of (0.5ug/mL) anti-CCL2 (R&D Systems, AF-479-SP). CD8^+^ T cells were isolated from the spleens of C57BL/6 mice using the MojoSort™ Mouse CD8^+^ T Cell Isolation Kit (Biolegend, #480035), followed by 1X Red Blood Cell lysis buffer (eBioscience, #00-4333-57) for 5 minutes. A 6.5mm insert with 8.0 μm polycarbonate membranes (COSTAR, #3422) was placed on top of the wells containing culture media. 100 μl of serum-free medium containing 2 × 10^5^ CD8^+^ T cells either with or without 24-hour treatment of (75ug/mL) anti-CXCR3 (Bio Cell, BE0249) were added to the top of the insert. Cells were incubated for 24 hours and then counted using a hemocytometer.

### *Ex vivo* co-culture of CD8^+^ T cells and cancer cells

YTN5 and YTN16 cells were transfected with FOXM1 siRNA for 24 hours and then exposed to 20ng/mL IFNγ (Abcam, #9922) for 24 hours. YTN5 and YTN16 cells were then trypsinized (Gibco, #15400-054) and seeded at 1 x 10^5^ in a 24 well plate and then pulsed with 1ng/mL OVA peptide (Sigma, #S7951-01mg) for 3 hours. Simultaneously, spleens were removed from C57BL/6-Tg (TcraTcrb) mice and splenocytes were treated with 1X Red Blood Cell lysis buffer (eBioscience, #00-4333-57) for 5 minutes. CD8^+^ T Cells were isolated using the MojoSort™ Mouse CD8^+^ T Cell Isolation Kit (Biolegend, #480035) and resuspended in RPMI-1640 medium (Corning, 45000-396) containing 20% FBS (Omega Scientific, FB-02). CD8^+^ T cells were incubated with 100U/mL IL2 (Abcam, #ab9856) for 24 hours, followed by 300ng/mL OVA peptide for 24 hours. CD8^+^ T cells were spun down at 1,500 x g for 5 minutes and resuspended at 4 x 10^5^ cells/100uL of RPMI-1640 medium (Corning, 45000-396) containing 20% FBS (Omega Scientific, FB-02). Media from YTN5 and YTN16 cells was removed and replaced with 400uL of fresh culture media, followed by 100uL of CD8^+^ T cell containing media. The co-cultured cells were incubated for 24 hours. The media and suspended CD8^+^ T cells were removed, and alive cancer cells were stained with 1% crystal violet as described above.

### ELISA

YTN5 and YTN16 cells were grown in 6 well plates and transfected with FOXM1 siRNA for 24 hours. Cells were then exposed to 20ng/mL IFNγ (Abcam, #9922) in 1 mL total volume for 24 hours. The confluence of cells was equal at the end of the treatment with IFNγ, and the media was collected for analysis. ELISA kits for CCL2 (R&D Systems, # DY479-05), CXCL9 (R&D Systems, # DY492-05), and CXCL10 (R&D Systems, # DY466-05) were performed according to the manufacturer’s protocols, using 100uL of media.

### Immunofluorescence Staining

EAC organoid cultures were fixed in 4% paraformaldehyde (Electron Microscopy Sciences, #15710) for 30 min. After washing organoids in PBS for three times, embedded in 2% agarose, dehydrated for making Paraffin blocks, and sectioned into 5-μm slices. Slides were deparaffinized and rehydrated, followed by antigen retrieval in boiling 10 mM sodium citrate buffer (pH 6.0) for 10 min. Slides were permeabilized in 0.3% Triton X-100 in PBS and blocked in 1% BSA in PBS for 30 min at room temperature. After blocking, slides were incubated with anti-Ki67 1:1000 (Abcam, #16667) for 3 hours in a humidified chamber at room temperature. Sections were washed by PBS with 0.1% Tween 20 (PBST) (three times for 5 min each) and incubated with Alexa Fluor secondary antibodies 1:500 (Thermo Fisher, #A-11036) for 1 hour. After washing with PBST, slides were mounted with Fluoroshield with DAPI (Sigma, #SLCD7376). Images were acquired with Keyence BZ-X810 microscopy.

### GSEA analysis of candidate TFs identified by ELMER

Based on the EAC tumor-specific DMRs that we identified recently, the ELMER method was applied to identify candidate transcription-factor-binding sequences (TFBS) and the top 30 TFs with q-value < 0.05 and average FPKM > 5 in EAC tumors were retained (10). For each candidate TF, we identified the nearest genes to the tumor-specific-hypoDMRs that contained the corresponding TFBS and ATAC-Seq peaks (from TCGA EAC samples) as the TF target genes (16). Next, GSEA was performed in PreRank mode using the fold change between EAC tumor and nonmalignant tissue as input and TF target genes as the library.

### RNA-seq data analysis

Paired end reads were aligned to the human reference genome (hg19) using STAR (– alignIntronMin 20–alignIntronMax 1000000 –alignSJoverhangMin 8 –quantMode GeneCounts) method. Differential gene expression analysis was performed using DESeq2 R package. RNA-Seq data for the OE19 cell line were downloaded from (E-MTAB-8579) and processed in a similar manner (44).

### ChIP-Seq data analysis

ChIP-Seq data of FOXM1 was generated in ESO26 and SKGT4 cell lines. Briefly, sequencing reads were aligned to human reference genome (HG19) using Bowtie2 (v2.2.6) (k = 2) (45). We used Picard MarkDuplicates tool to mark PCR duplicates. ENCODE blacklisted regions were removed. Macs2 was utilized to identify the peaks with the parameters –bdg –SPMR –nomodel –extsize 200 –q 0.01. Bigwig files were generated by bamCompare in DeepTools (v3.1.3) using parameters –binSize 10– numberOfProcessors 5 –scaleFactorsMethod None –normalizeUsing CPM– ignoreDuplicates –extendReads 200 from Ramirez et al.,2014 (46). The bigwig files were visualized in Integrative Genomics Viewer (IGV) (47).

### Data availability

The mRNA expression (RNA-Seq level-3 data) data of EAC patient samples were retrieved from either GSE1420, MTAB4504, or TCGA (GDC v16.0) using TCGAbiolinks (V2.14.1) R package. ChIP-Seq data of FOXM1 generated from the OE33 cell line were from (22). H3K27Ac ChIP-Seq data in the ESO26 cell line was generated by us previously (21). ATAC-seq and RNA-seq upon ERBB2-knockdown were from (44). The blacklisted regions were downloaded from ENCODE database (https://sites.google.com/site/anshulkundaje/projects/blacklists). Molecular Signatures Database v7.4 (https://www.gsea-msigdb.org/gsea/msigdb/index.jsp) were used for GSEA (v3.0) (https://www.gsea-msigdb.org/gsea/index.jsp). The ChIP-Seq and RNA-Seq generated in this study have been deposited in the Gene Expression Omnibus (GEO) repository (GSE236847, token-mvghwgaqljyptyd). The remaining data are available in the article or Supplementary Information files, or available from the authors upon request. The full scans of Western blotting and the data presented in a plot, chart or other visual representation format were provided in the Source Data file. Source data are provided with this paper.

## Acknowledgements

We would like to thank Sachiyo Nomura for kindly gifting the YTN cells.

## Author Contributions

D.-C.L. conceived and devised the study. D.-C.L., B.Z., Q.Y., M.S., Y.Z., C.N., and H.Z., designed experiments and analyses. B.Z., Q. Y., M.S., Y. Z., C. N., and H. Z. performed the experiments. Q.Y. and Y.Z. performed bioinformatics and statistical analysis. U.K.S. contributed reagents and materials. B.Z., Q. Y., M.S., Y. Z., C. N., and H. Z., analyzed the data. D.-C.L. and U.K.S. supervised the research. B.Z. and D.-C.L. wrote the manuscript.

## Ethics Approval

Patient samples were approved by the Institutional Review Board, IRB HS-22-122. All animal experiments were approved by Institutional Animal Care and Use Committee (IACUC) of USC, Protocol-21383.

