## Supplemental Figures for "Epigenomic Analyses Identify FOXM1 As a Key Regulator of Anti-Tumor Immune Response in Esophageal Adenocarcinoma"

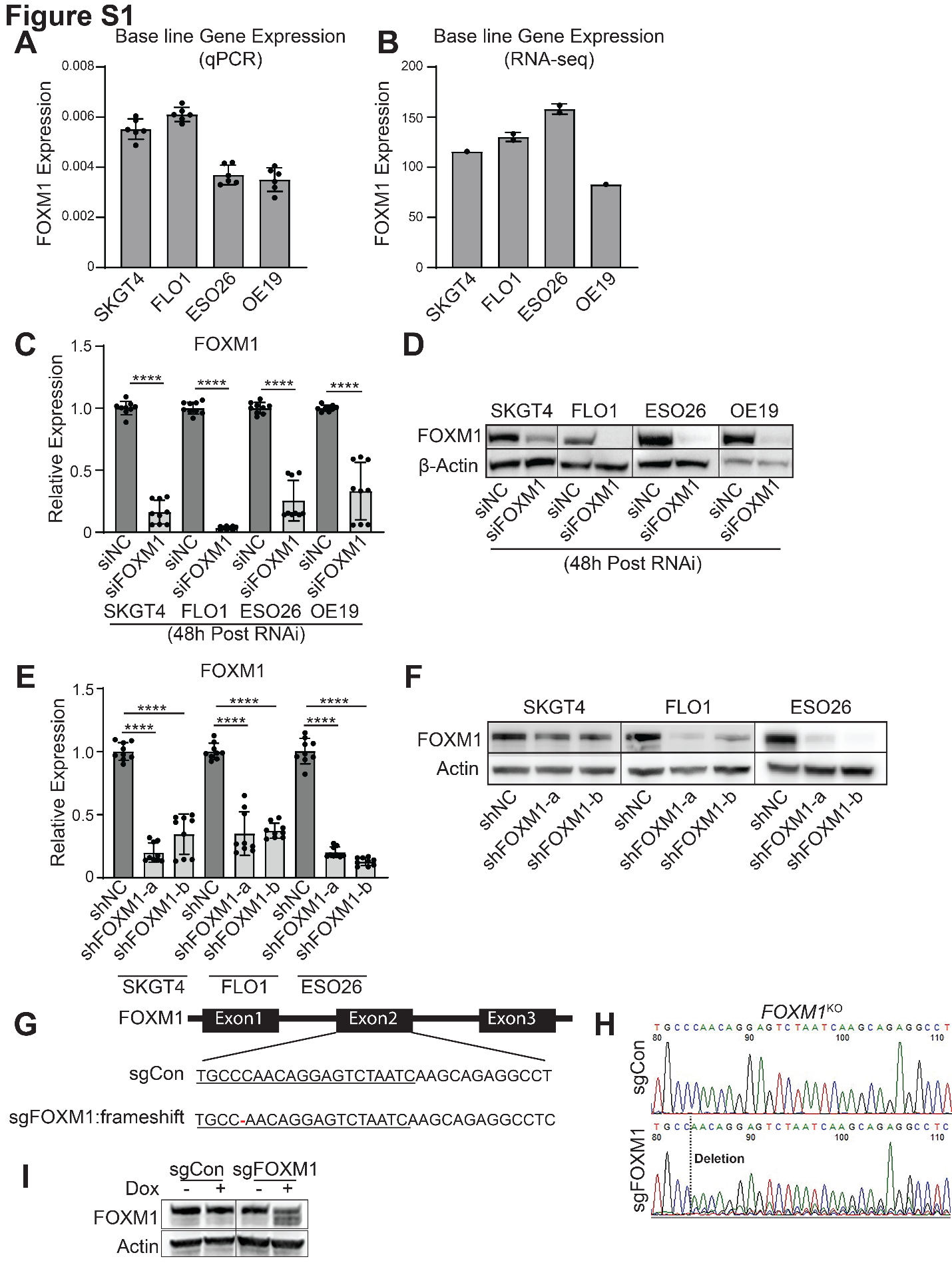
**Figure S1.** Bar graphs showing the baseline gene expression of FOXM1 in EAC cells using (**A**) qPCR and (**B**) RNA-seq data. Bars are shown as the means ± SD for 2 replicates and as the mean for 1 replicate. (**C**) Bar graph showing the relative expression of FOXM1 upon 48-hour treatment with siRNA for FOXM1. Shown are the means ± SD from three technical replicates. ****P<0.0001; P-values were determined using an ordinary one-way ANOVA. (**D**) Western blot detection of FOXM1 in EAC cell lines upon 48-hour treatment with siRNA for FOXM1. Blot shown is representative of one replicate. (**E**) Bar graphs showing relative expression of FOXM1 upon FOXM1 knockdown cells with two different shRNA targets. Shown are the means ± SD from three technical replicates. ****P<0.0001; P-values were determined using an ordinary one-way ANOVA. (**F**) Western blot detection of FOXM1 in EAC cell lines upon FOXM1 knockdown cells with two different shRNA targets. Blot shown is representative of two replicates. (**G**) Approach for targeting FOXM1 using CRISPR Cas9. A guide RNA specific for exon 2 of FOXM1 was designed to create a deletion of a single base, causing a frameshift mutation. **(H**) Sanger sequencing showing the effects of the CRISPR Cas9 single base deletion for the guide RNA upon induction using doxycycline in the EAC cell line SKGT4. (**I**) Western blot detection of FOXM1 in the EAC cell line SKGT4 upon induction with doxycycline. Blot shown is representative of one replicate.

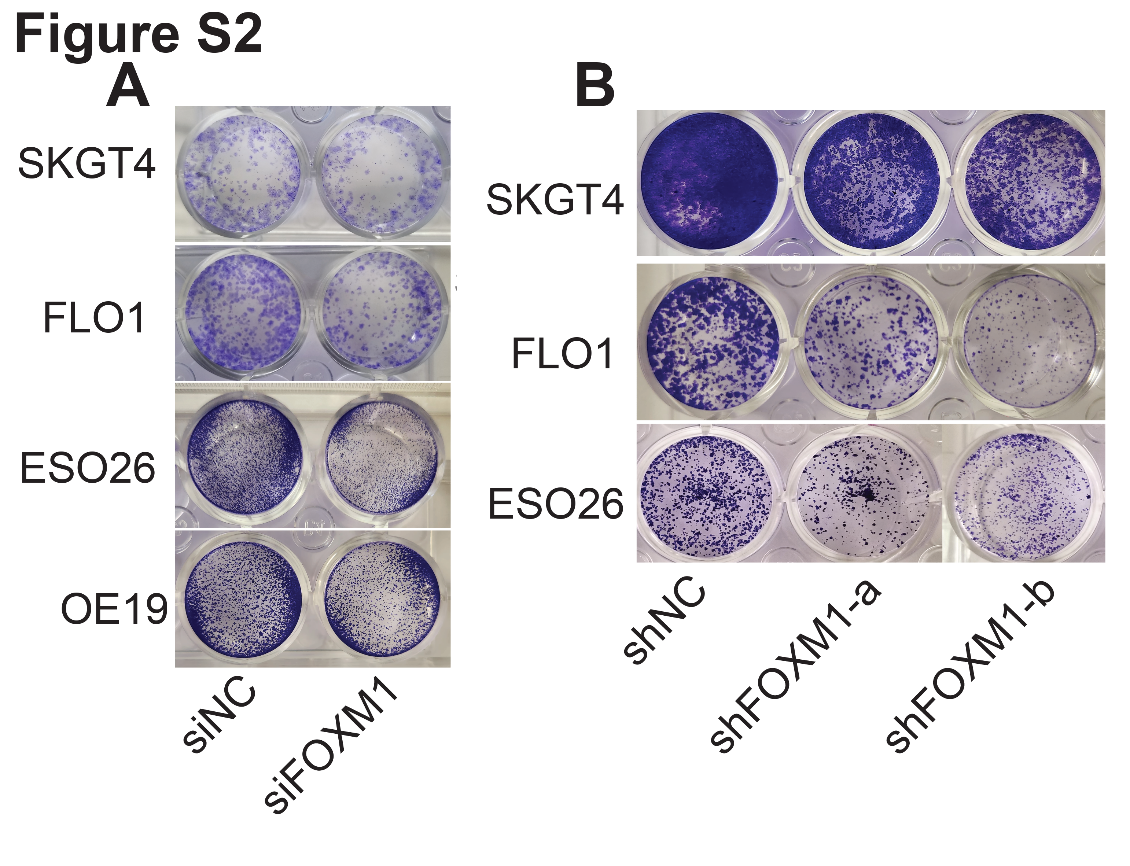

**Figure S2.** Representative images of colony formation upon (**A**) treatment with siRNA for FOXM1 and (**B**) FOXM1 knockdown cells with two different shRNA targets. Images shown are representative of three replicates.
